# The assembled Banana dihaploid mitochondrial genome is compact with a high number of gene copies

**DOI:** 10.1101/2022.01.13.476214

**Authors:** Shruthy Priya Prakash, Vaidheki Chandrasekar, Selvi Subramanian, Rahamatthunnisha Ummar

**Affiliations:** PSG College of Technology, Department of Biotechnology, Coimbatore, Tamil Nadu, India 641004

**Keywords:** Mitochondrial genome, *Musa acuminata*, DH Pahang, genome assembly, RNA editing, mitochondrial genes

## Abstract

Banana being a major food crop all around the world, attracts various research interests in crop improvement. In banana, complete genome sequences of *Musa accuminata* and *Musa balbisiana* are available. However, the mitochondrial genome is not sequenced or assembled. Mitochondrial (mt) genes play an important role in flower and seed development and in Cytoplasmic Male Sterility. Unraveling banana mt genome architecture will be a foundation for understanding inheritance of traits and their evolution. In this study, the complete banana mt genome is assembled from the whole genome sequence data of the *Musa acuminata subsp. malaccensis* DH-Pahang. The mt genome sequence acquired by this approach was 409574 bp and it contains, 54 genes coding for 25 respiratory complex proteins 15 ribosomal proteins, 12 tRNA genes and two ribosomal RNA gene. Except atpB, rps11 and rps19 other genes are in multiple copies. The copy number is 12 in tRNA genes. In addition, nearly 25% tandem repeats are also present in it. These mt proteins are identical to the mt proteins present in the other members of AA genome and share 98% sequence similarity with *M. balbisiana.* The C to U RNA editing is profoundly higher (87 vs 13%) in transcripts of *M. balbisiana* (BB) compared to *M. accuminata* (AA). The banana AA mitochondrial genome is tightly packed with 233 genes, with less rearrangements and just 5.3% chloroplast DNA in it. The maintenance of high copy number of functional mt genes suggest that they have a crucial role in the evolution of banana.

## INTRODUCTION

The genomes of present day cultivated bananas consist of either diploids, triploids or tetraploids of AA (*Musa acuminata*), BB (*M. balbisiana*) and combination of both (AB). The mitochondrial (mt) genome of banana is not sequenced/assembled to date. The available AA genomic (D’Hont et al 2012) and BB genomic (Wang et al. 2019) sequence resources help to reconstitute the mt genomes. NOVOPlasty is a *denovo* organellar genome assembly tool which uses Whole Genome Shotgun (WGS) sequences to assemble circular organellar genomes (Dierckxsens et al. 2017). This program was tested to assemble the mt genomes of model plants rice and Arabidopsis and gave 99.9% accuracy. Norgal (de Novo ORGAneLle extractor) is one another pipeline available to extract the organellar DNA from the Whole genome Shotgun (WGS) sequences (Al-Nakheeb et al. 2017). Using this, full circular mt genome of Panda, a sea weed, butterfly and fungal genomes were assembled with 99.5 % sequence similarity with reference sequences. Wang et al., (2018) used Newbler, Amos and Minimus software to assemble mt and chloroplast genomes of an ornamental plant *Salix suchowensis,* and a fruit tree, Ziziphus jujuba. In the above methods there is no need to separately isolate the organellar DNA of high quality. *Brassica oleracea* var. capitata mt genome assembly from WGS is found to be 219,975 bp in size with no large repeats (Yang et al.2018). Recently, mt genomes of *Sinapis arvensis var.* ‘Yeyou 18’, a cytoplasmic male sterile line (Nsa CMS) and its corresponding maintainer line ‘Zhongshuang 4’were assembled using the mt sequences present in the total DNA (Sang et al. 2020). Present work deals with *Insilico* approaches to assemble and reconstitute the mt genome of banana species *Musa accuminata* using the WGS data in comparison with other model plants such as maize, rice, sugar cane, sugar beet and Arabidopsis.

The mt genome size of land plants range between 66 kb in *Viscum scurruloidem* to 11.3Mbp in *Siline conica* (Omelchenko et al. 2020). Among them the herbaceous monocots have a range of 400-500 kb (Cuenca et al. 2013). Plant mt genomes evolve in a dramatic burst due to the presence of several repeats (Wynn and Christensen 2019). The reasons for the larger mt genome size in plants and lesser number of assembled mt genomes compared to animals are also due to the rearrangements and recombinations caused by these repeats (Kovar et al. 2018). Comparison of mt genome sizes of a diploid (A_2_) and an allotetraploid (AD_2_) cotton species indicated only a slight variation in size and found to be 644 and 677 kb respectively (Chen et al. 2017). The cotton A, D and AD mt genomes differed drastically and had four or six large repeats leading to lot of inversions and translocations. Mt genomes of rice wide crosses and backcross inbred lines have shown radical change in the gene order and copy number (Yang et al. 2020). In Fabales, a different kind of genome expansion has occurred in mt genomes. Horizontal gene transfer events between intercellular and interspecific level have led to the genome size variation (Choi et al. 2019). Considerable amount of mt DNA in plants are horizontally transferred to either nuclear or chloroplast genome. DNA transfer events from mitochondria to nucleus are positively correlated to the size of nuclear genomes in several plants (Zang et al. 2020). Often, these horizontally transferred genes are not activated (Pinard et al.2019).

The mt gene content in plants do not vary much. Barley wild and cultivated varieties were found to have similar mt genome size and gene content with only three SNPs. These genomes contain 33 protein coding genes, three rRNA and 16 tRNAs (Hisano et al. 2016). Similarly, chiltepin pepper *(Capsicum annuum* var *glabriusculum)* has 31 known protein coding genes, three rRNA genes and 25 tRNA genes (Magdy and Ouyong 2020). In *Raphanus sativus,* L. 40 protein coding genes, three rRNA genes and 23 tRNA genes were found in the mt genome (Peng and Gao 2020). Very recently, six mitogenomes of *Damnacanthus indicus* was sequenced and found to have 32 protein coding genes after several losses (Han et al. 2021). Plant organellar genes undergo post transcriptional modifications such as splicing and editing, among them RNA editing specifically C to U change is found to be evolved during early land plant development (Liu et al. 2011). Most of the editing sites create nonsynonymous changes leading to protein change however they lead to a conservative change to maintain the function (Omelchenko et al. 2020). RNA editing in protein coding genes is found to increase the protein function and codon bias. Higher frequency of editing was observed immediately after exposure to salt stress in Barley mt nad3 gene (Ramadan 2020). However, in Arabidopsis reduced RNA editing rate is found in heat stressed plants and suggested to have a regulatory role in abiotic stress tolerance (Chu and Wei 2020). Differential RNA editing pattern was observed between Cytoplasmic Male Sterile (CMS) and Fertile plants in Pigeon Pea (Kaila et al. 2019). Nodulation process which fixes atmospheric Nitrogen is an energy demanding development involving higher mt activity. Higher splicing and RNA editing efficiency is seen in nodulating roots (Sun et al. 2020). Edera and Sanchez-Puerta (2021) has recently developed a computational frame work to identify editing sites in *Nicotiana tabacum* mt genome.

Assembly of AA mt genome will provide the genome size, number of respiratory and ribosomal proteins, tRNA and rRNA genes in it. Assembling the organellar genome from the whole genome sequence data can be possibly done de novo by using programs like Norgal (Al-Nakeeb et al. 2017) or by using reference sequence with CONTIGuator (Galardini et al.2015, Halim et al.2016). This information will be useful for researchers who are interested in analyzing characters which are controlled by nuclear and mitochondrial interaction (Hanson and Bentolila 2004). In Maize (Weiwei et al. 2017) and Arabidopsis (Lee et al. 2017) embryo, seed development is modulated by genes located in mt genome which is regulated by nuclear genes. Mitochondrial genes undergo post transcriptional modifications such as splicing and editing which are regulated tissue and stage specifically by nuclear genes (Hanson and Bentolila 2004). The knowledge on gene content, gene structure cis or trans spliced would enable a researcher to correlate a molecular factor to a phenotype. This study focuses on assembling the mt genome of banana for the above-mentioned applications. Besides, paternal vs maternal transmission (Faure et al.,1994) of mt genes to the hybrids can be ascertained if the full genome is known in completion.

## METHODOLOGY

### Data collection

The contigs using WGS of the *Musa acuminata* subsp*. malaccensis* was collected from NCBI GenBank CAJGYN000000000.1 (Bioproject: PRJEA82777) (D’Hont et al. 2012). Among the genomic sequences,12 mt fragments have been separated from the nuclear genome. The contigs were quality checked removed from the other contigs and stored in a separate file. These contigs were then subjected to nucleotide search using NCBI BLASTN using 5 different reference mt genomes of Maize (NC_007982.1), rice (NC_011033), Arabidopsis (NC_037304.1), sugar-beet (NC_002511.2) and sugarcane (NC_031164.1).

### Sequence assembly and circularization

The sequence comparison of the mt contigs of *Musa acuminata* subsp. *malaccensis* DH Pahang resulted in several fragments of sequence from each reference. The sequences obtained as a result of NCBI BLASTN are pooled together and assembled using CONTIGuator (Galardini et al.2015) (http://contiguator.sourceforge.net/) that resulted in a single scaffold. This program uses BLAST to align the draft sequence against the reference sequence and provides a single scaffold. CGView (Grant and Stothard 2008) (http://cgview.ca/) is a Circular Genome Visualization server was used to depict a circular genome of mitochondria. The scaffold sequence (1.2 Mb) as well as assembled sequence in single fasta format were analyzed through CG viewer. Gene labeling in the map was done in the same tool. Individual track of protein coding genes, ORFs, tRNAs and rRNAs were created as separate text file and fed to the server for visualization. GC skew and GC content can be obtained from the server directly. The location of the genes whether in sense or antisense strand were also manually recorded while labeling. Legend and captions were also included in the figure through the software.

### Repeats Identification

Repeats in the assembled mitochondrial genome was identified using Tandem Repeat Finder which is a public repository tool to identify repeats present in genomic DNA. (https://tandem.bu.edu/cgi-bin/trdb/trdb.exe). The assembled mt genome in fasta file format was fed to the server to find repeats. Result was immediately obtained with coordinates, sequences and copy number. It can be downloaded in any formats for further analyses. The copy number and percentage of repeats were calculated using the obtained results.

### Gene Annotation

MFANNOT is a mt genome annotation (https://megasun.bch.umontreal.ca/RNAweasel/) (Beck and Lang 2010) server that utilizes various tools and programs to provide detailed annotation on the introns, exon sequences using comparative analysis. The input given is the scaffold sequence obtained from the CONTIGuator, as well as the merged fasta file and the parameters are set to default with standard mitochondrion annotation settings. Time taken for annotation depends on the size of the mt genome given as input, as the plant mt genomes are bigger in size, total time needed can be up to 1-2 hours. The results are directly sent to the email entered to the server. Results consists of 3 files, one with the sequences translated, one with the annotations and other with the genes present. The GC content of the scaffold was calculated using GC calculator. The translated sequences are separated and stored for future use. Annotated information is manually recorded in MS excel sheet for easy handling of information. The results were also compared with another annotation tool MITOFY (https://dogma.ccbb.utexas.edu/mitofy/).

### Comparative Analyses and Editing site identification

The annotated list of mt genes identified in *Musa accuminata subsp. malaccensis* DH-Pahang are compared against the available genomic data of other varieties, subspecies, species and genus by doing a BLASTP search. The results were analyzed for variation in gene content, split genes and length variation in nucleotide and protein sequences with reference to the *M. balbisiana* mt genes. Chloroplast DNA content in banana mt genome was analyzed by BLASTN search of both mt and chloroplast genomes of banana. Editing sites for all the mt genes of *M. accuminata* and *M. balbisiana* were identified using their DNA compared against their respective transcriptome data (SRX10460839; SRX767394) available in NCBI. Pairwise comparison of mt DNA and transcript sequences (cDNA) were carried using BLASTN program for all annotated genes. Variations corresponding to C in mt DNA and T in cDNA were considered as C to U edited sites.

## RESULTS

### Mitochondrial genome assembly

As a result of banana mt sequences assembly using the CONTIGuator, with five model organisms as references, a single scaffold was obtained. The scaffold sequence was 409,574 bp in length with a GC content of 45.3% (Table 1). The percentage coverage and identity obtained in comparison with related model organisms, Maize, rice, sugarcane, sugar beet and Arabidopsis are given in Fig 1. Annotation of this assembled genome by MFANNOT resulted in the identification 97 genes including ORFs (Table 1; Fig. 2). There are 45 ORFs present in the genome which are 300bp and above reported by the program. The number of respiratory genes in banana mt genome is 25, ribosomal protein genes 15, tRNA genes 12 and rRNA gene one. The coding regions of the genome accounts to 22.8% of the total genome. There are 169 tandem repeats present in the banana mt genome (Supplementary Table 1). The length of the repeats ranges from 15 bp to 1500 bp. These tandem repeats accounts to about 25.34% of the total mt genome size. The two copy repeats contribute to 21.22%. The three and four copy repeats contribute to 2.61% and 1.51% of the mt genome size respectively.

**Fig 1:**
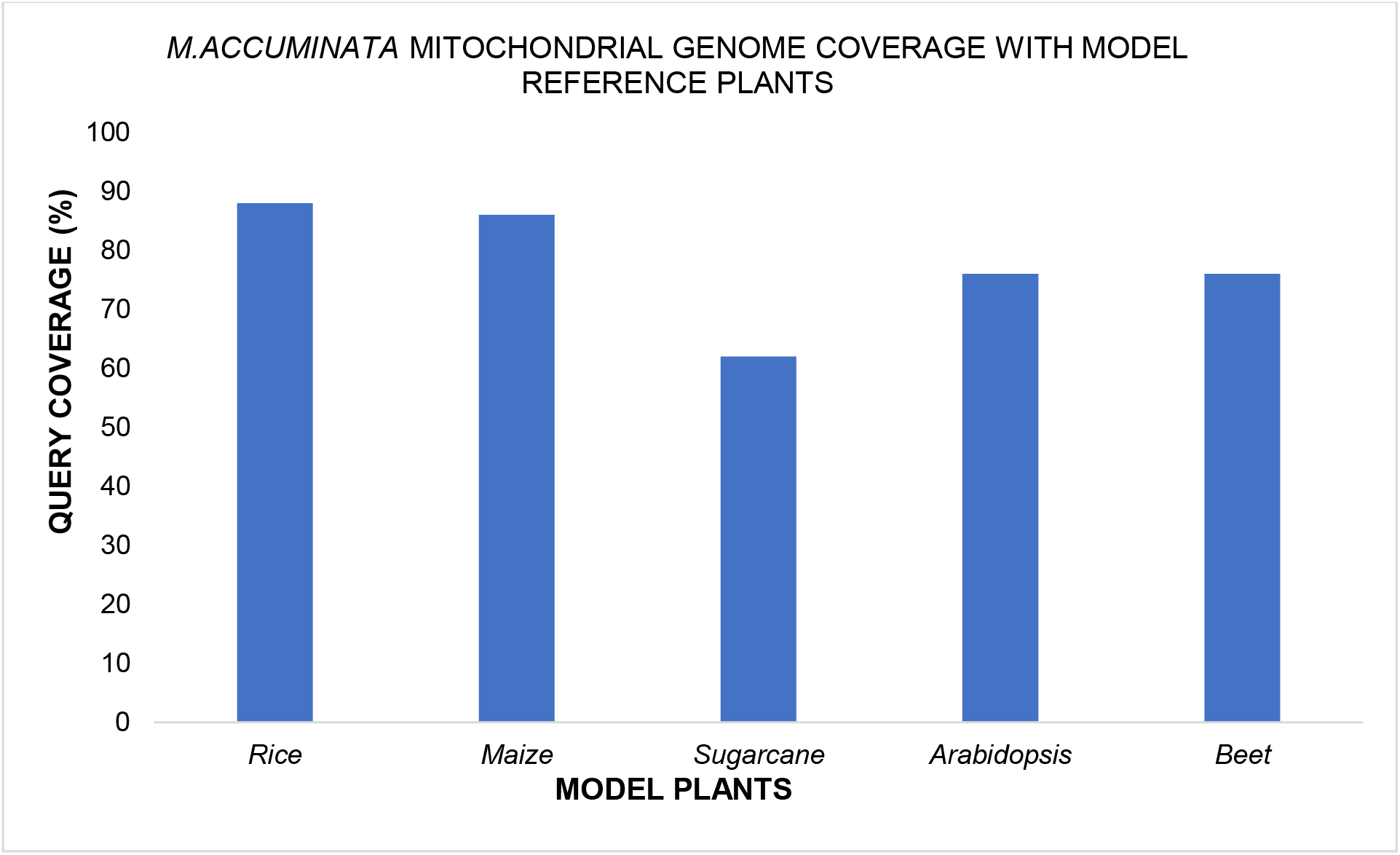
Graph showing query coverage of *M.accuminata* mitochondrial genome with Rice, Maize, Sugarcane, Arabidopsis and Beet mitochondrial genomes

**Table 1:** General characters of the assembled *M. accuminata* mitochondrial genome.

| Features | Content |
| --- | --- |
| Genome size | 409754 bp |
| GC content | 45.3% |
| Chloroplast content | 5.3% |
| Tandem repeats | 25.34% |
| Respiratory complex | 22 |
| Ribosomal protein | 10 |
| Other protein | 1 |
| Pseudo genes | 7 |
| tRNAs | 12 |
| rRNA | 2 |
| ORF | 43 |
| Total no of genes | 97 |

### Mitochondrial gene Analyses

The location, copy number, presence and number of introns of these genes are listed in the Table 2. The number of genes with known unique function excluding the additional copies is 41 (Table 2). Out of these 41 genes only three of them are single copy genes. Other genes have maximum of seven copies. The number of copies of tRNA is found to be even more and one of them have fourteen copies. (Supplementary table 2). There are ten split genes present in the mt genome. These genes also contain groupII introns. Among them, atpB and ccmF are cis-spliced and the rest eight of them are trans-spliced genes. Among the cis-spliced genes, atpB has two introns, and ccmF is with single intron in four copies and one copy has two introns. Among the 233 genes including all copies present in the mt genome (Fig. 2) 141 of them are present in sense strand and the rest in antisense strand. A comparative analyses of gene content with the four model organisms, maize, rice, Arabidopsis and sugarbeet are presented in Table 3. Among the respiratory complex genes, ccmFC and ccmB genes are not present in banana with reference to the four model organisms. Ribosomal protein genes, rps1, rps7 and rpl2 are pseudogenes. Other category genes, matr, mttB are absent compared to the other organism whereas ftsH is a pseudogene in banana. There are seven pseudogenes identified in the annotation results which are all truncated in nature. However, there are few genes which are intact with unconventional start and stop codons (Table 2). The amino acid length and the percentage similarity of the mt proteins of other *M. accuminata* subspecies or varieties show 100% similarity except a few (supplementary Table 2). The average percentage similarity shared between *M. accuminata* and *M. balbisiana* mt proteins is 98% (Table 2). There are four protein coding genes that are missing in *M. balbisiana* genome compared with *M.accuminata.*

**Fig 2:**
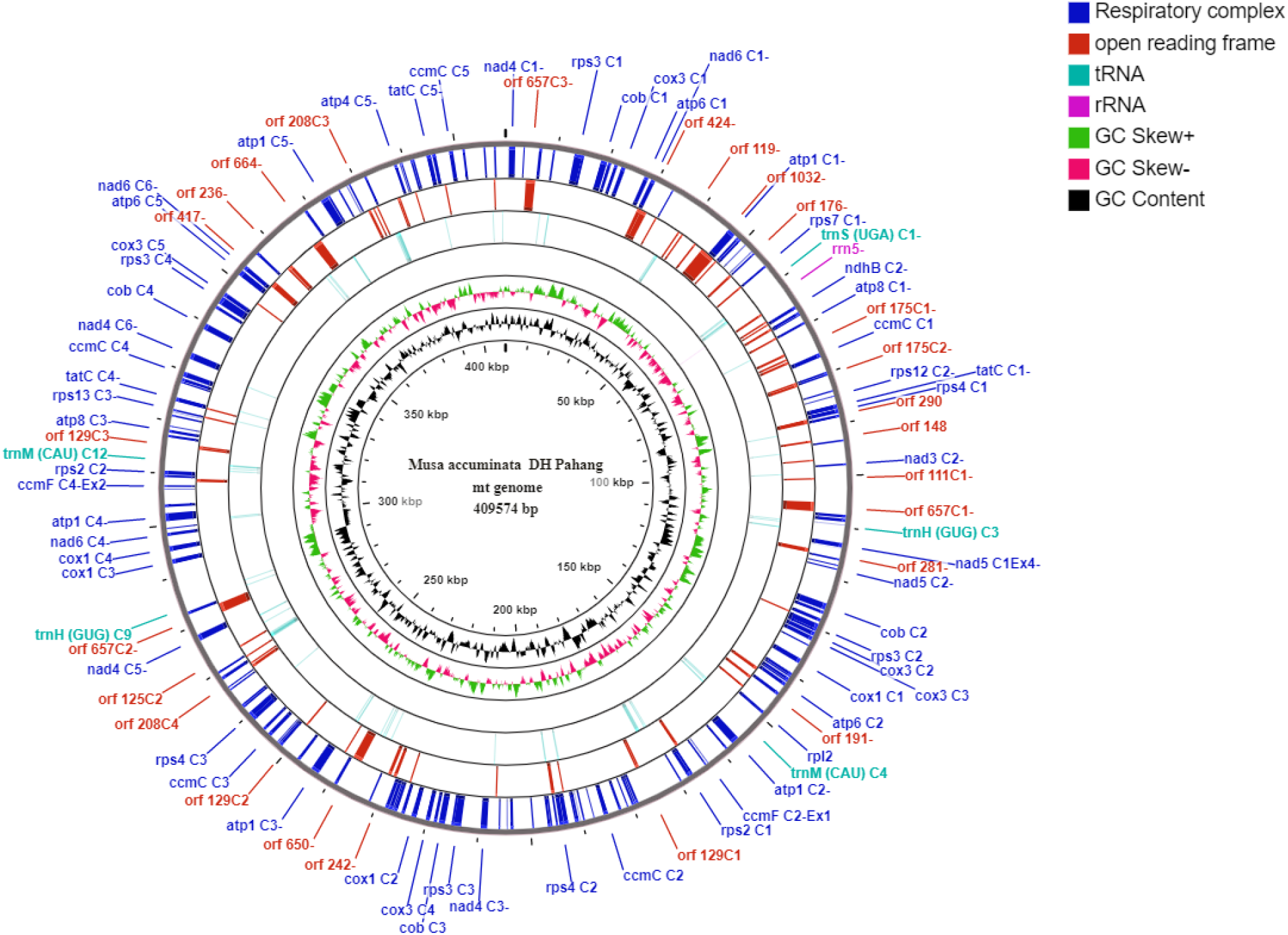
Circular view of assembled *M.accuminata* subsp. *malaccensis* DH Pahang with individual tracks which depicts respiratory complex, open reading frame, tRNA and rRNA genes, along with GC skew and GC content

**Table 2:**
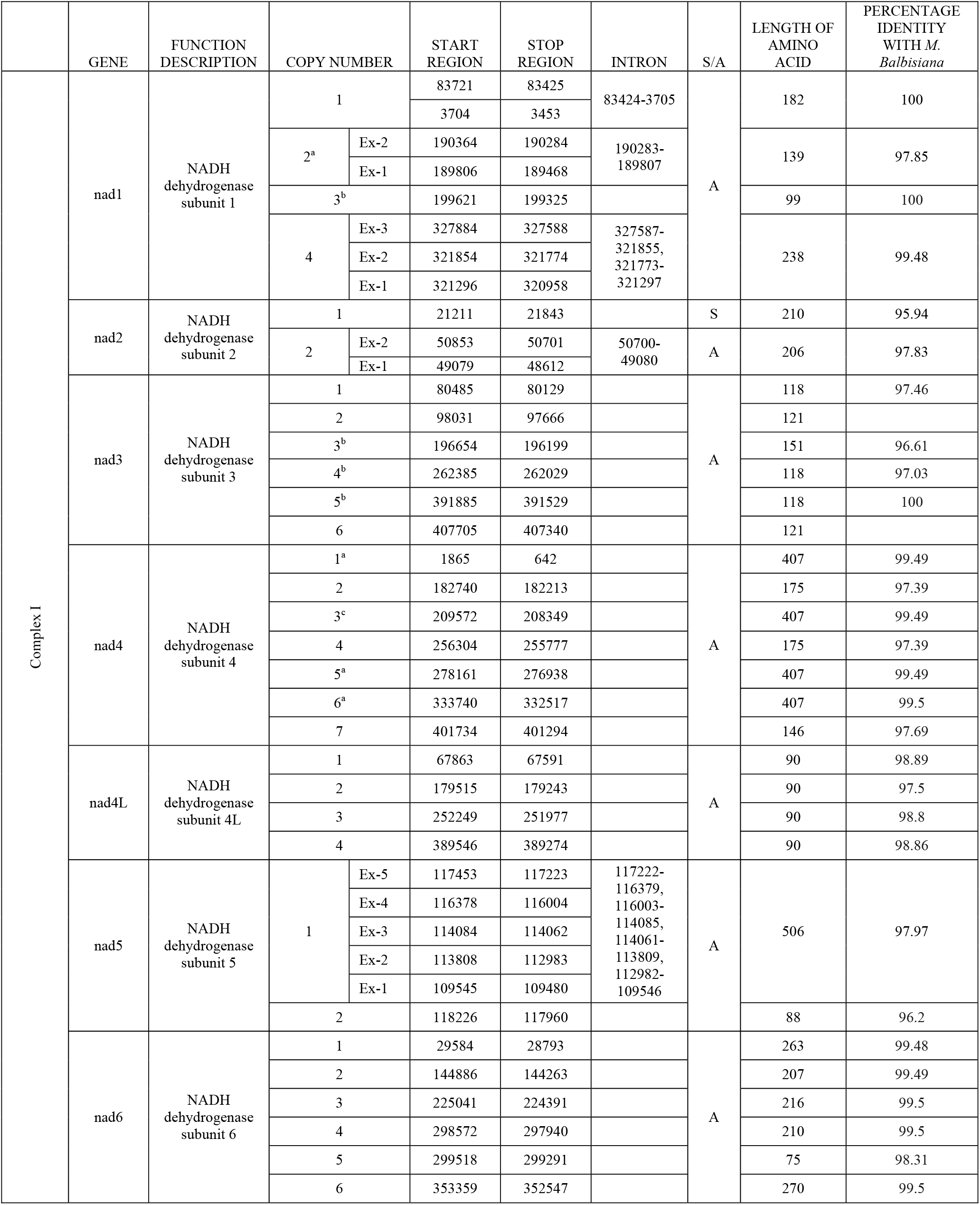
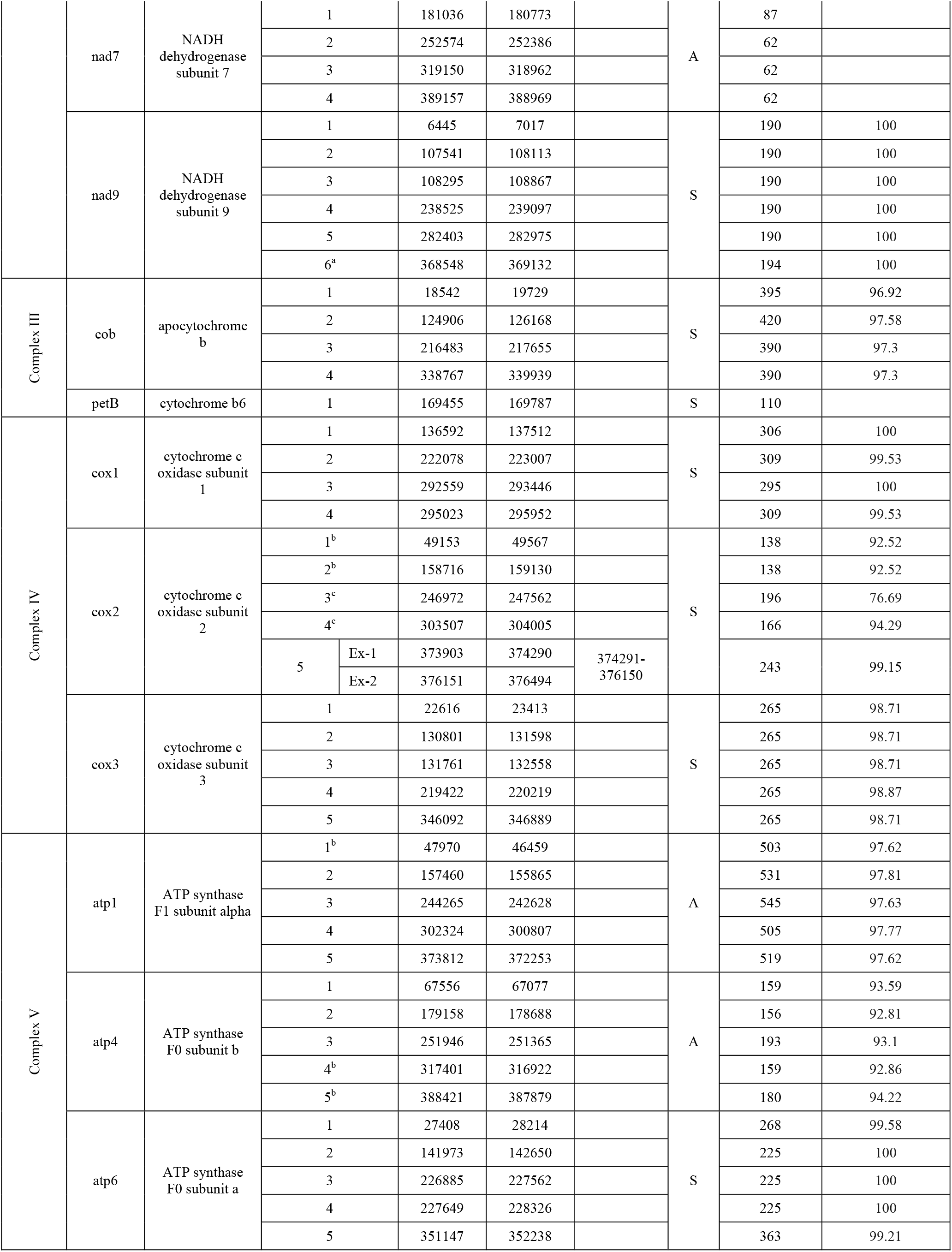
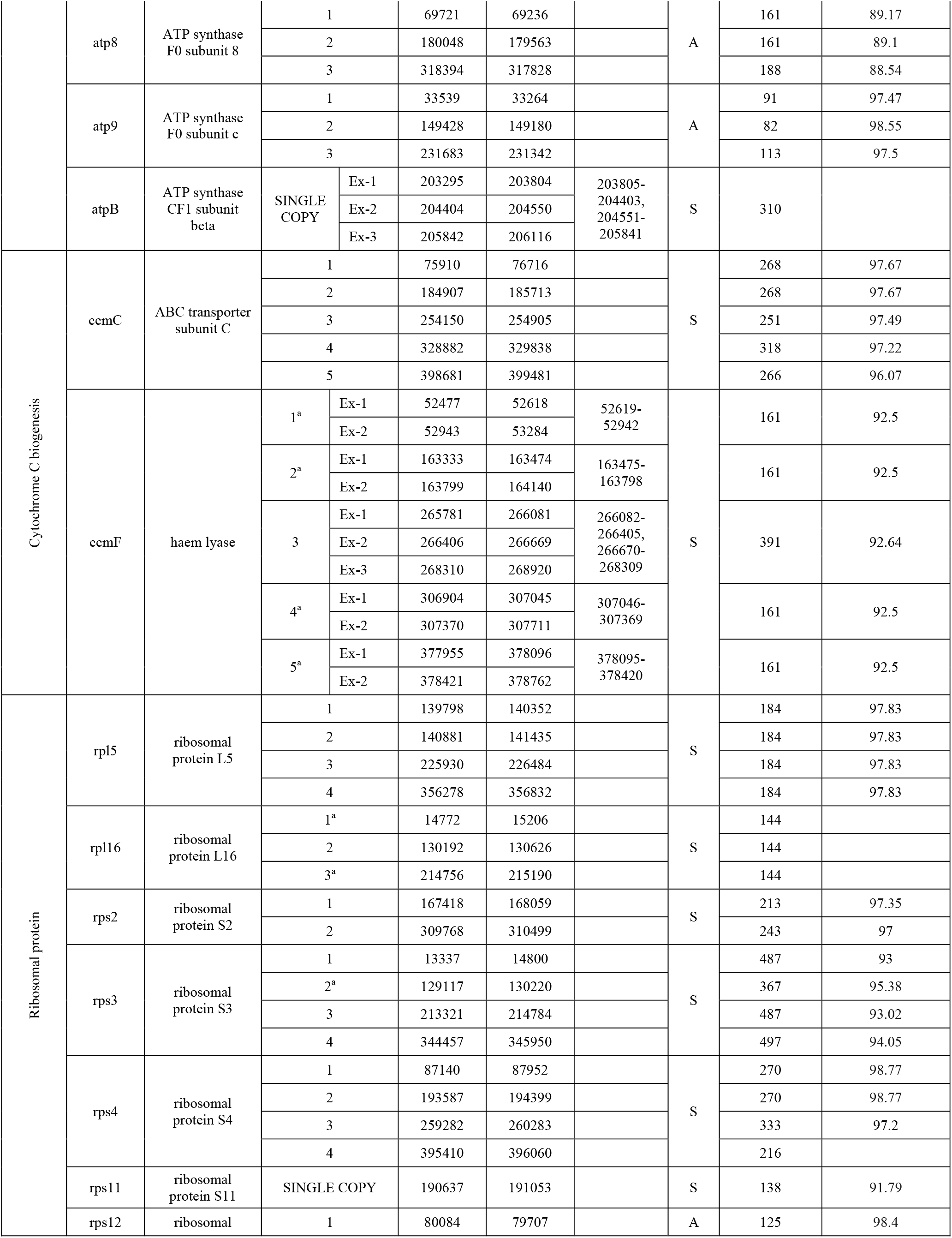
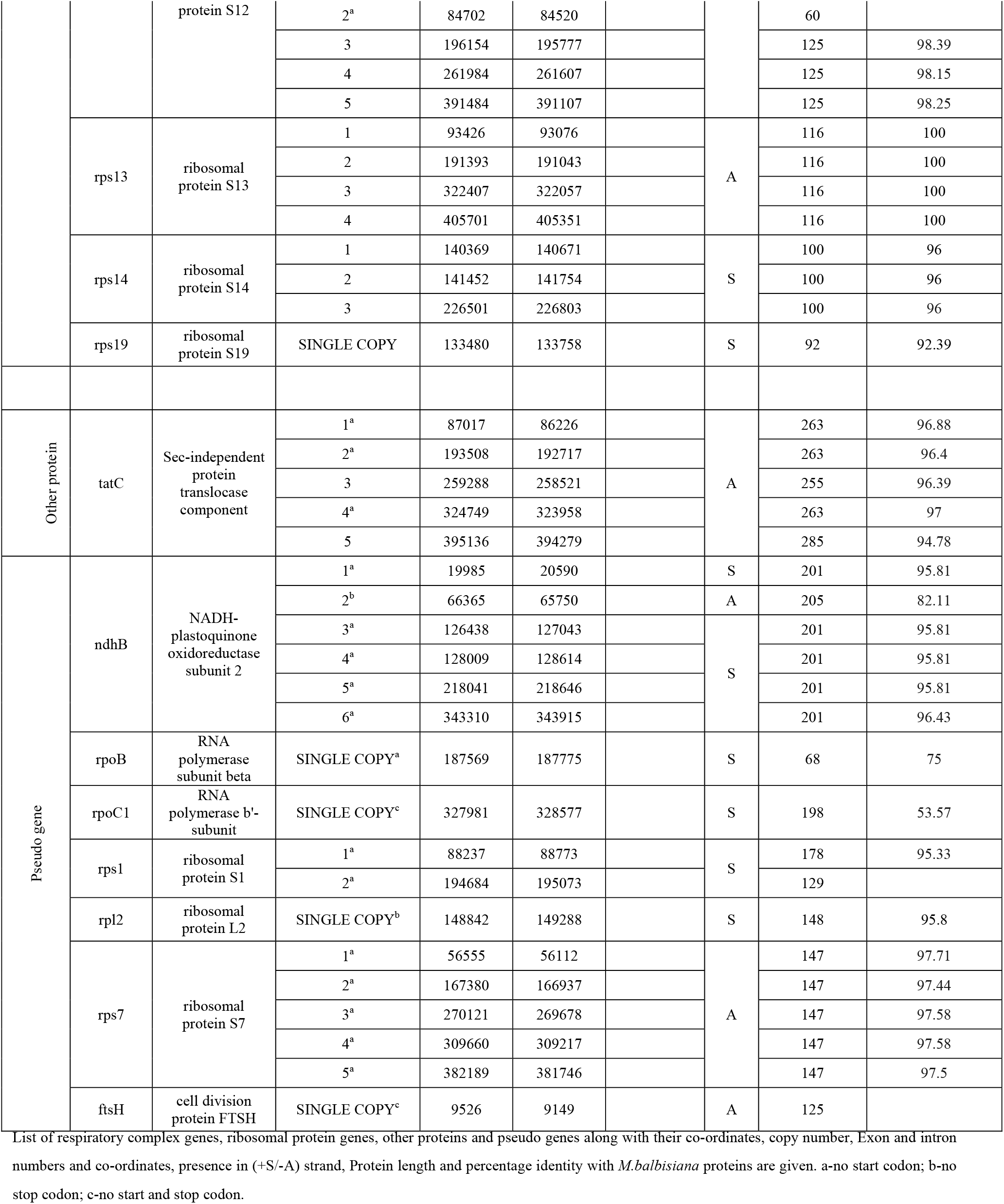
List of mitochondrial genes annotated in *M.accuminata* DH Pahang subspecies malaccensis and their comparison with *M. balbisiana*.

**Table 3:**
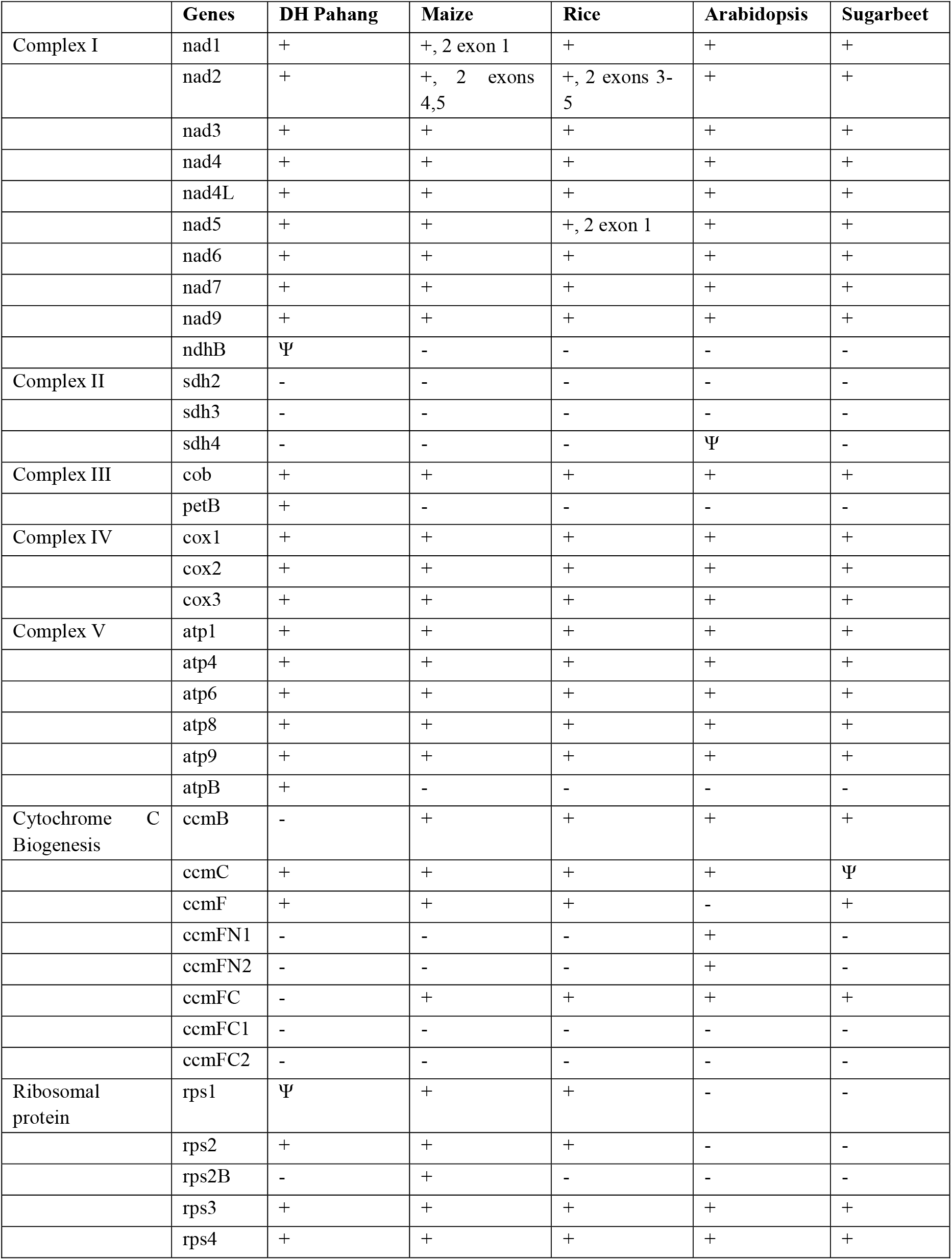
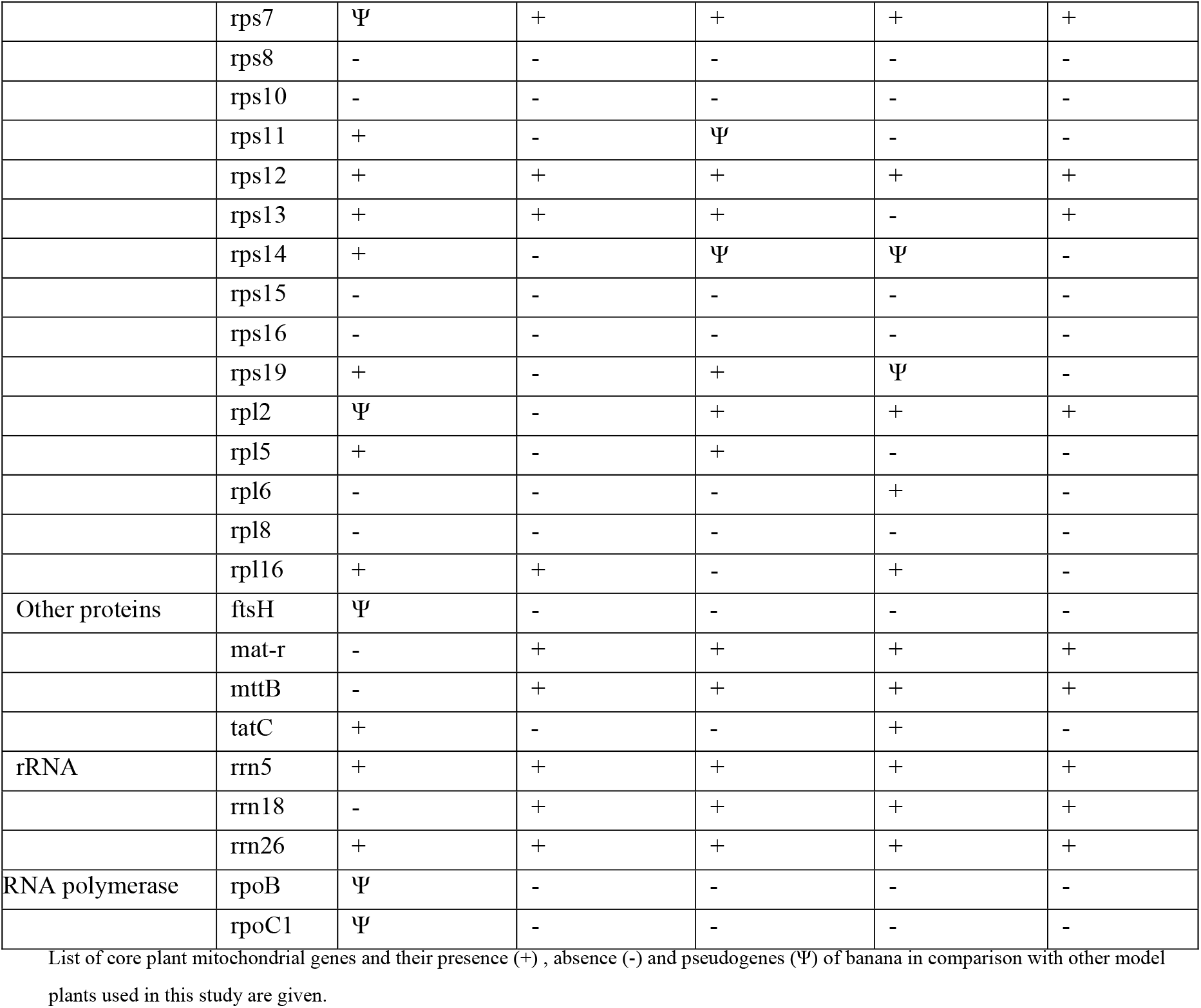
*M.accuminata* mitochondrial gene content comparison with other model organisms used as reference in this study.

### Mitochondrial RNA editing

The mt RNA editing status was analyzed for all protein coding genes of *M. accuminata* and compared against the *M. balbisiana.* The number of editing sites, their position, change of amino acid if any and intron position are given in table 4. There is a drastic difference (41 vs 6) observed in the editing status between *M. balbisiana* and *M. accuminata* mt genomes. The graph (Fig. 3) represents the genes in which editing differences are found between *M. accuminata* and *M. balbisiana*. The bar length corresponds to the number of editing sites observed. There were 18 genes which showed editing of mRNA in *M. balbisiana* species compared to only five genes in *M. accuminata*. Among them nad6 and rps1 are unique which were not edited in *M. balbisiana.* The other three genes, ccmC, rps12 and petB genes had just one editing site. The nad6 gene is the only gene that had two editing sites in Accuminata however one of them is a synonymous change (Table 4). Twenty seven percent of the editing observed in *M. balbisiana* are synonymous. atp 9 gene had the maximum of seven editing sites and a stop codon is created by editing. Introns are also found to be edited in nad4L and nad5 genes of Balbisiana.

**Fig 3:**
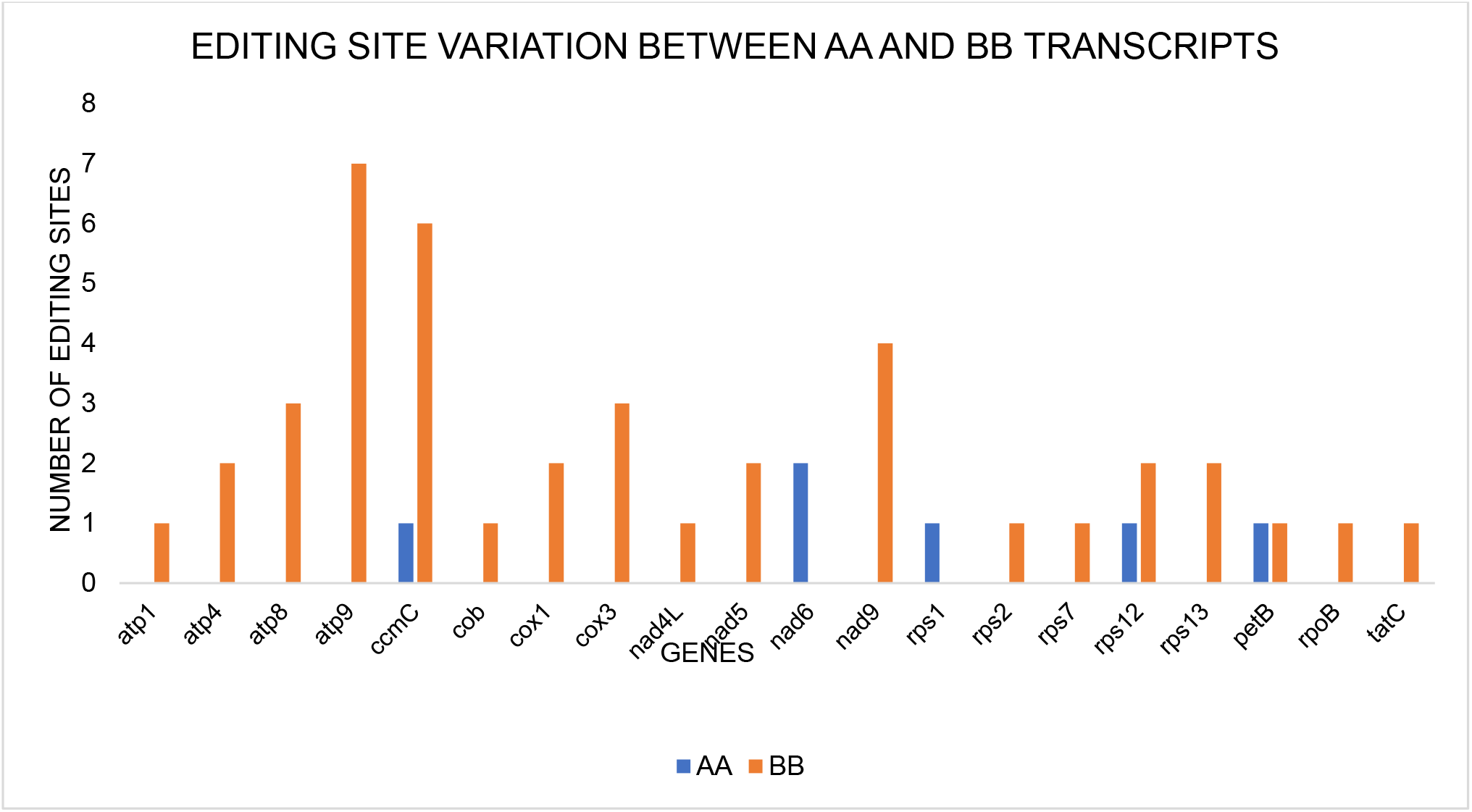
Bar graph showing number of C to U editing sites identified in mitochondrial transcripts between M.accuminata (blue) and M.balbisiana (orange). Length of the bar represents the number of editing sites in respective genes

**Table 4:**
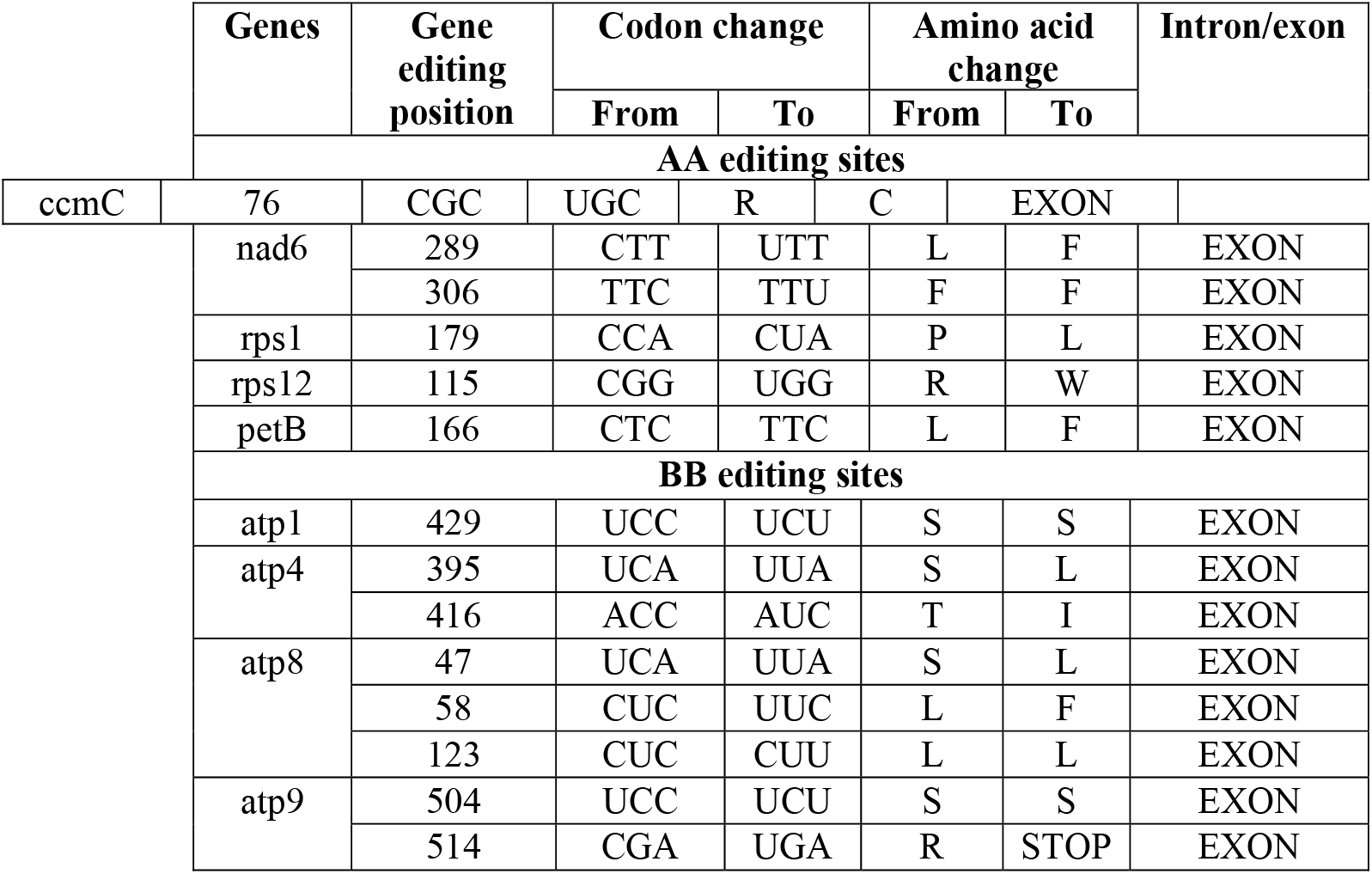
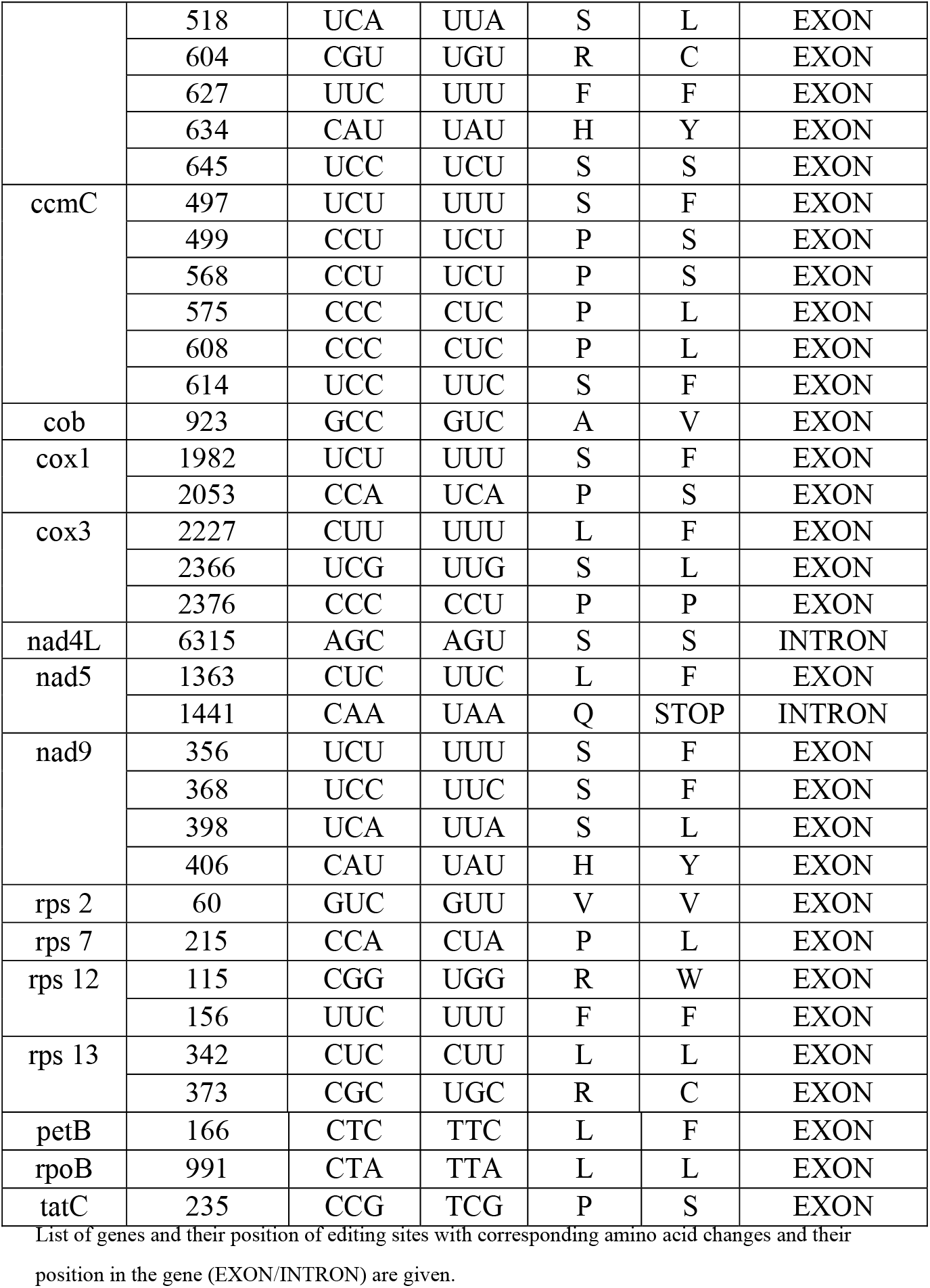
List of editing sites found in*M.accuminata* transcripts in comparison with*M.balbisiana*.

|  | Genes | Gene editing position | Codon change |  | Amino acid change |  | Intron/exon |
| --- | --- | --- | --- | --- | --- | --- | --- |
|  |  |  | From | To | From | To |  |
| AA editing sites |  |  |  |  |  |  |  |
| ccmC | 76 | CGC | UGC | R | C | EXON |  |
|  | nad6 | 289 | CTT | UTT | L | F | EXON |
|  |  | 306 | TTC | TTU | F | F | EXON |
|  | rps1 | 179 | CCA | CUA | P | L | EXON |
|  | rps12 | 115 | CGG | UGG | R | W | EXON |
|  | petB | 166 | CTC | TTC | L | F | EXON |
|  | BB editing sites |  |  |  |  |  |  |
|  | atp1 | 429 | UCC | UCU | S | S | EXON |
|  | atp4 | 395 | UCA | UUA | S | L | EXON |
|  |  | 416 | ACC | AUC | T | I | EXON |
|  | atp8 | 47 | UCA | UUA | S | L | EXON |
|  |  | 58 | CUC | UUC | L | F | EXON |
|  |  | 123 | CUC | CUU | L | L | EXON |
|  | atp9 | 504 | UCC | UCU | S | S | EXON |
|  |  | 514 | CGA | UGA | R | STOP | EXON |

|  |  |  |  |  |  |  |
| --- | --- | --- | --- | --- | --- | --- |
|  | 518 | UCA | UUA | S | L | EXON |
|  | 604 | CGU | UGU | R | C | EXON |
|  | 627 | UUC | UUU | F | F | EXON |
|  | 634 | CAU | UAU | H | Y | EXON |
|  | 645 | UCC | UCU | S | S | EXON |
| ccmC | 497 | UCU | UUU | S | F | EXON |
|  | 499 | CCU | UCU | P | S | EXON |
|  | 568 | CCU | UCU | P | S | EXON |
|  | 575 | CCC | CUC | P | L | EXON |
|  | 608 | CCC | CUC | P | L | EXON |
|  | 614 | UCC | UUC | S | F | EXON |
| cob | 923 | GCC | GUC | A | V | EXON |
| cox1 | 1982 | UCU | UUU | S | F | EXON |
|  | 2053 | CCA | UCA | P | S | EXON |
| cox3 | 2227 | CUU | UUU | L | F | EXON |
|  | 2366 | UCG | UUG | S | L | EXON |
|  | 2376 | CCC | CCU | P | P | EXON |
| nad4L | 6315 | AGC | AGU | S | S | INTRON |
| nad5 | 1363 | CUC | UUC | L | F | EXON |
|  | 1441 | CAA | UAA | Q | STOP | INTRON |
| nad9 | 356 | UCU | UUU | S | F | EXON |
|  | 368 | UCC | UUC | S | F | EXON |
|  | 398 | UCA | UUA | S | L | EXON |
|  | 406 | CAU | UAU | H | Y | EXON |
| rps 2 | 60 | GUC | GUU | V | V | EXON |
| rps 7 | 215 | CCA | CUA | P | L | EXON |
| rps 12 | 115 | CGG | UGG | R | W | EXON |
|  | 156 | UUC | UUU | F | F | EXON |
| rps 13 | 342 | CUC | CUU | L | L | EXON |
|  | 373 | CGC | UGC | R | C | EXON |
| petB | 166 | CTC | TTC | L | F | EXON |
| rpoB | 991 | CTA | TTA | L | L | EXON |
| tatC | 235 | CCG | TCG | P | S | EXON |

List of genes and their position of editing sites with corresponding amino acid changes and their position in the gene (EXON/INTRON) are given.

## DISCUSSION

### Banana mt genome assembly from Whole Genome Shotgun sequence

The contigs of mt sequences of DH Pahang were assembled from the genomic shotgun sequences. This approach of extracting mt sequences from total genomic sequence was followed in several algal mt genome assembly. Mitochondrial genomes of diatoms *Phaeodaetylum tricornutum* and *Talassiosira pseudonana* were obtained by sequencing the total genome including plastid, mt and nuclear genomes (Secq and Green 2011). The shotgun sequences were assembled by JGI/JAZZ assembler. The mt genomes were assembled by sequence similarity to other algal mt genomes. Similarly, sequences of mt genomes of 10 algae were assembled from the genomic sequences downloaded either from genbank or related published data (Guillory et al.2018). These algal mt sequences were selected based on GC content, size, BLASTN sequence similarity to other mt genomes. Falcon (Ver.4) was used to assemble the genome and circular mt topology was predicted. CONTIGuator tool was used to assemble the mt sequences listed in the DH Pahang CAJGYN000000000.1. This tool can address the sequence gaps and more than one circular molecule (Galardini et al.2011). Besides, this can also align contigs from a draft genome by comparing it to several reference genomes based on their alignment and orientation (Galardini et al. 2015).

### Compact mt genome

The mt genome size of banana is 409 kb. This is smaller than two other monocots, maize, 539 kb (Clifton et al. 2004) and rice, 490 kb (Notsu et al. 2002) mt genomes. The smaller size implicates compactly arranged genes with lesser extent of repeated sequences. The gene number is found to be comparatively higher (97 vs 58) than maize mt genome. Similarly, when the number of tandem repeat sequences were analyzed most of them are two copies and the three and four copy number repeats contribute to only 4% of the genome (Supplementary Table 1). In sugar beet, the repeat sequence TR1, which contains an array of 32 bp repeat sequences was found to be repeated to a maximum of 13 times (Nishizawa et al. 2000). In solanaceous plants, a common origin of these short repeats was observed. However, only in members of the tribe Hyoscyameae it has expanded to eight copies (Gandhini et al. 2019). Melon mt genome which is 2740 kb has nearly 42% of repeat sequences (Rodrigues – Moreno et al. 2011). These repeat sequences beside contributing to the mt genome size, also involve in recombination events that can further increase the genome size. Sullivan et al. (2020) have performed comparative analysis on the repeat abundance and recombination frequency in plant mt genomes. They reported that the recombination dynamics was heterogenous among gymnosperms and short repeats of 200bp and below are actively involved in recombination in one third of plant species they analyzed. There is a possibility of at least one recombination event in banana mt genome also.

Nearly 40% of the genes are in the antisense strand and the rest in the sense strand, there are only three shifts in orientation in the full genome (Table 2 and Fig 2.). Presence of promiscuous DNA from other compartments by horizontal transfer might also contribute to the size of the mt genomes in plants. In apple approximately 20% of the mt DNA is transferred from other compartments (Goremykin et al. 2012). In banana, just 5.3% of chloroplast DNA was found in the mt genome (Table 1). The above observations, small genome size, possibility of one or two recombination events, high gene copies and low chloroplast DNA content could be the reasons for the compact nature of the genome.

### High gene copy number

Most of the banana mitochondrial genes are found duplicated and the maximum copy number is six in many of these genes. This suggests genome duplication rather than gene duplication event. Segmental duplication of genes can be identified when looking at the gene order repeated (Fig. 2). For example, *rps12-nad3-nad1* order is repeated in the banana mt genome. *M. accuminata* genome is a dihaploid with AA genome. The process of dihaploid development by tissue culture may have increased the copy number of genes in this banana mt genome. Furthermore, paternal transmission and the possibility of maintaining several subcircular forms of the intact mt genome may contribute to this high copy number (Nagata et al., 1999; Tsujimura et al. 2019). In yeast, a linear scale increase in copy number with ploidy number is observed in both genomes (De Chiara et al. 2020). Similarly, in Arabidopsis autoployploids, nuclear mitochondrial coordination is observed in genome duplication followed by tuned gene expression patterns in mt genes (Coate et al. 2020). The copy number of tRNA genes are higher than the protein coding genes. Out of the 12 tRNA genes five of them match to the chloroplast genes. Chloroplast tRNA genes are found functional in mitochondria of angiosperms (Richarson et al. 2013). In Magnolia six tRNA genes are transferred from plastid and they are functional (Dong et al. 2020). This suggests that some of the tRNA could have chloroplast origin in banana also. RNA mediated gene duplication events are found in several taxa of plants (Cuenca et al. 2012). They have observed existence of processed paralogues along with precursors of nad1 genes. In the case of banana DH Pahang mt genome, DNA duplication is most likely as introns are retained in all copies. However, the possibility of precursor mRNA/cDNA insertion is also likely since additional introns are gained in ccmF gene. Intact nature of these additional copies of these genes in banana suggests that they are under functional constraint.

### Gene content

The total number of unique protein coding genes present in the banana mitochondrial genome is slightly higher than other model plants. The core protein coding genes observed in Angiosperms is 24 (reviewed by Zardoya 2020). Mangnolia has the largest set of 64 unique mitochondrial genes among flowering plant (Dong et al. 2020). However, the largest mt (11.7 Mb) genome assembled to date is from a gymnosperm which has 77 unique genes (Putintseva et al. 2020). The total number of unique genes is 52 in banana which goes up to 233 when all copies are considered. Hence the genome is compact with very little intergenic space. The number of tRNA and rRNA genes are slightly lower than other mt genomes. The average number of tRNA genes found in mt genomes across angiosperms is 17-29 (Michaud et al. 2011). In banana, the copy number of tRNA genes is up to twelve. The number of pseudogenes is seven in banana mt genome; many of them could be remnant genes that might have been recently transferred to nucleus (Subramanian et al. 2001). The rpl2 gene is one example, which is a pseudogene in wheat, functional in rice whereas many other plants do not have a mitochondrial copy. In banana, also a truncated rpl2 is present, the rest of the gene could not be matched, but the annotation result suggests both possibility of pseudogene and a trans-spliced gene. A complete analysis of these truncated pseudogenes in closely related organisms in both nuclear and mt compartment might provide evidence for the recent horizontal transfer events. Recently, O’conner and Li (2020) also have proposed ‘Mitochondrial Fostering’ theory in which mitochondria play an important role in arrival and development of orphan genes which are not present in any other plants. In banana, there are several genes with either or both start and stop codon missing or non-classical are present. The unconventional start and stop codons may not make the gene to pseudogene. In Arabidopsis ccmF (N2) gene is functional despite the absence of classical start codon (Rayapuram et al. 2008). The tatC is a membrane transport protein which is present other than respiratory complex and ribosomal proteins in banana. This is also found in few other species including moss Lycopodium cernuum (Kanagara et al. 2021).

### Banana A and B mitochondrial genome comparison

Comparative analyses of *M. accuminata* sub sp *malacensis* DH-Pahang mt proteins with other AA genome mt proteins show 100 percentage similarity (Supplementary Table 3). In some multicopy genes, one copy of these genes shares a slightly lesser similarity however that is higher than the similarity percentage found between the two species *M. Accuminata* and *M. balbisiana*. The average sequence similarity between these two Musa species is 98%. Barley and wheat mitochondrial respiratory proteins were almost similar with only few differences in few genes (Hisano et al. 2016). In cotton allopolyploids, among A and D genomes, A is found to be the donor of mitochondrial genomes of the progenitors (Chen et al. 2017). In banana also A and B genomes are available, comparing B mitochondrial genome and autopolyploid progenitors would resolve the origin of banana mt genome. In fishes, the mitochondrial gene similarity between allopolyploids were found to be higher when transmitted maternally than paternal transmission (You et al. 2014). The paternal/biparental transmission of mitochondrial DNA is documented in banana (Nagata et al.1999). When the editing sites were compared between two Musa genomes, Balbisiana is having a drastic high number. However, the editing sites were not leading to conservative changes in B genome. Similarly, A genome editing sites were different to that of B genome editing event. The A and B genome sequences used for comparison are from wild genotypes that are under natural selection. Hence, they may be evolving independently.

## Data availability

The data underlying this article are available in NCBI (National Center for Biotechnology Information) and can be accessed with accession numbers -CAJGYN000000000.1-*Musa acuminata* subsp*. malaccensis,* NC_007982.1-Maize, NC_011033-rice, NC_037304.1-Arabidopsis, NC_002511.2-sugar-beet and NC_031164.1-sugarcane. The assembled AA genome data will be submitted to NCBI after annotation.

## Acknowledgement

We thank all the authors and fellows who supported this idea and their encouragement towards completing this article.

## Funding

This study was initiated as part of a “North East Banana Consortium funded project, by Department of Biotechnology, Government of India. It is a full in silico work, only the second Authors salary was from the fund.

## Conflicts of interest/Competing interests

No conflicting interest

